# AVADA Enables Automated Genetic Variant Curation Directly from the Full Text Literature

**DOI:** 10.1101/461269

**Authors:** Johannes Birgmeier, Andrew P. Tierno, Peter D. Stenson, Cole A. Deisseroth, Karthik A. Jagadeesh, David N. Cooper, Jonathan A. Bernstein, Maximilian Haeussler, Gill Bejerano

## Abstract

**Purpose:** The primary literature on human genetic diseases includes descriptions of pathogenic variants that are essential for clinical diagnosis. Variant databases such as ClinVar and HGMD collect pathogenic variants by manual curation. We aimed to automatically construct a freely accessible database of pathogenic variants directly from full-text articles about genetic disease.

**Methods:** AVADA (Automatically curated VAriant DAtabase) is a novel machine learning tool that uses natural language processing to automatically identify pathogenic variants and genes in full text of primary literature and converts them to genomic coordinates for rapid downstream use.

**Results:** AVADA automatically curated almost 60% of pathogenic variants deposited in HGMD, a 4.4-fold improvement over the current state of the art in automated variant extraction. AVADA also contains more than 60,000 pathogenic variants that are in HGMD, but not in ClinVar. In a cohort of 245 diagnosed patients, AVADA correctly annotated 38 previously described diagnostic variants, compared to 43 using HGMD, 20 using ClinVar and only 13 (wholly subsumed by AVADA and ClinVar’s) using the best automated abstracts-only based approach.

**Conclusion:** AVADA is the first machine learning tool that automatically curates a variants database directly from full text literature. AVADA is available upon publication at http://bejerano.stanford.edu/AVADA.

## Introduction

Rare genetic diseases affect 7 million infants born every year worldwide^1^. Exome or genome sequencing is now entering clinical practice in aid of the identification of molecular causes of highly penetrant genetic diseases, and in particular Mendelian disorders (genetic diseases caused by pathogenic variants in a single gene^2–4^). In a Mendelian context, typically one or two of the patient’s genetic variants in a single gene are causative of the patient’s disease. After following standard variant filtering procedures, a typical singleton patient exome contains 200-500 rare functional variants^5^. Identifying causative variants is therefore very time-consuming, as investigating each variant and deciding whether or not it is causative can take up to an hour^6^. Various approaches are in development to accelerate this process^7–10^. Identifying causative variants can be greatly accelerated if the patient’s genome contains a previously reported pathogenic variant that partly or fully explains their phenotype. The American College of Medical Genetics (ACMG) guidelines for the interpretation of sequence variants recommend variant annotation using databases of reported pathogenic variants^11^.

The rapidly growing literature on human genetic diseases^12^, the costly process of manual variant curation^13^, and improved computational access to the full text of primary literature^14,15^ serve to incentivize automatic variant curation. Creating a variant database from the primary literature involves finding variant descriptions (such as “c.123A>G”), linking them to a transcript of the correct gene mention, and converting them to genomic coordinates (chromosome, position, reference and alternative alleles) so they can be readily intersected with any patient variants. Previous work on automatic variant discovery in the literature has largely focused on finding variant descriptions in paper titles and abstracts with high accuracy without converting the discovered variants to genomic coordinates^16–22^. Previous automatic variant curation tools have also focused on mapping variant mentions to dbSNP^23^ variant identifiers (rsIDs). Mapping textual variant descriptions directly to reference genome coordinates requires significant effort, and has thus far largely been left to manually curated databases such as HGMD^24^ and ClinVar^25^, which devote many thousands of wo/man-hours to the task of collecting genetic variants from either the scientific literature or clinical laboratories.

The recently started ClinGen project has proposed to “develop machine-learning algorithms to improve the throughput of variant interpretation”^26^ and note that a rate limiting factor for clinical use of variant information is the lack of openly accessible knowledgebases capturing genetic variants. We posed the question as to whether manual variant curation to genome coordinates can be accelerated with the help of machine learning approaches by first training an automatic curation system on a sample of manually curated variants (from ClinVar and HGMD), and then applying the trained system to the entire body of PubMed indexed literature for automatic curation of published variants. AVADA (Automatically curated VAriant DAtabase), our automated variant extractor, identifies variants in genetic disease literature and converts all detected variants into a database of genomic (GRCh37/hg19) coordinates, reference and alternative alleles. We show that AVADA improves on the state of the art in automated variant extraction, by comparing it to tmVar 2.0^27^, a best-in-class tool used to harvest variants from PubMed abstracts. Combining the free ClinVar and AVADA variant databases, we find that we can recover a significant fraction of diagnostic disease-causing variants in a cohort of 245 patients with Mendelian diseases.

## Materials and Methods

### Identification of relevant literature

PubMed is a database containing titles and abstracts of biomedical articles, only a subset of which contain descriptions of variants that cause human genetic disease. A document classifier is a machine learning classifier that takes as its input arbitrary text and classifies it as “positive” (here, meaning an article about genetic disease) or “negative” (otherwise). We trained a scikit-learn^28^ LogisticRegression^29^ classifier to identify relevant documents using positive input texts (titles and abstracts of articles cited in OMIM^30^ and HGMD^24^) and negative input texts (random titles and abstracts from PubMed). Machine learning classifiers take as input a real-valued vector (the “feature vector”) describing the input numerically. Input texts were converted into a feature vector by means of a scikit-learn CountVectorizer followed by a TF-IDF^31^ transformer (an operation that converts input text to a feature vector based on the frequency of words in input documents). After training the title/abstract document classifier, we applied it to all 25,793,020 titles and abstracts in PubMed to identify articles that might be relevant to the diagnosis of genetic diseases. Full text PDFs of relevant articles were then downloaded and converted to text using pdftotext^32^ version 0.26.5. Because identifying potentially relevant articles based upon title and abstract alone often yields articles whose full text does not turn out to be relevant for the diagnosis of genetic diseases, we subsequently trained a full-text scikit-learn LogisticRegression classifier to classify downloaded full-text documents as “relevant” or “irrelevant” based upon the article’s full text. As with the title/abstract classifier, full text documents were converted to a feature vector by means of a CountVectorizer followed by a TF-IDF transformer. Filtering full-text articles for relevance resulted in a subset of downloaded articles more relevant to the diagnosis of genetic disease (Supplementary Methods). A total of 133,410 articles were downloaded and subsequently classified as relevant to the diagnosis of human genetic diseases based on the articles’ full text. We refer to this set of articles as the “AVADA full-text articles” (Figure 1).

**Figure 1.**
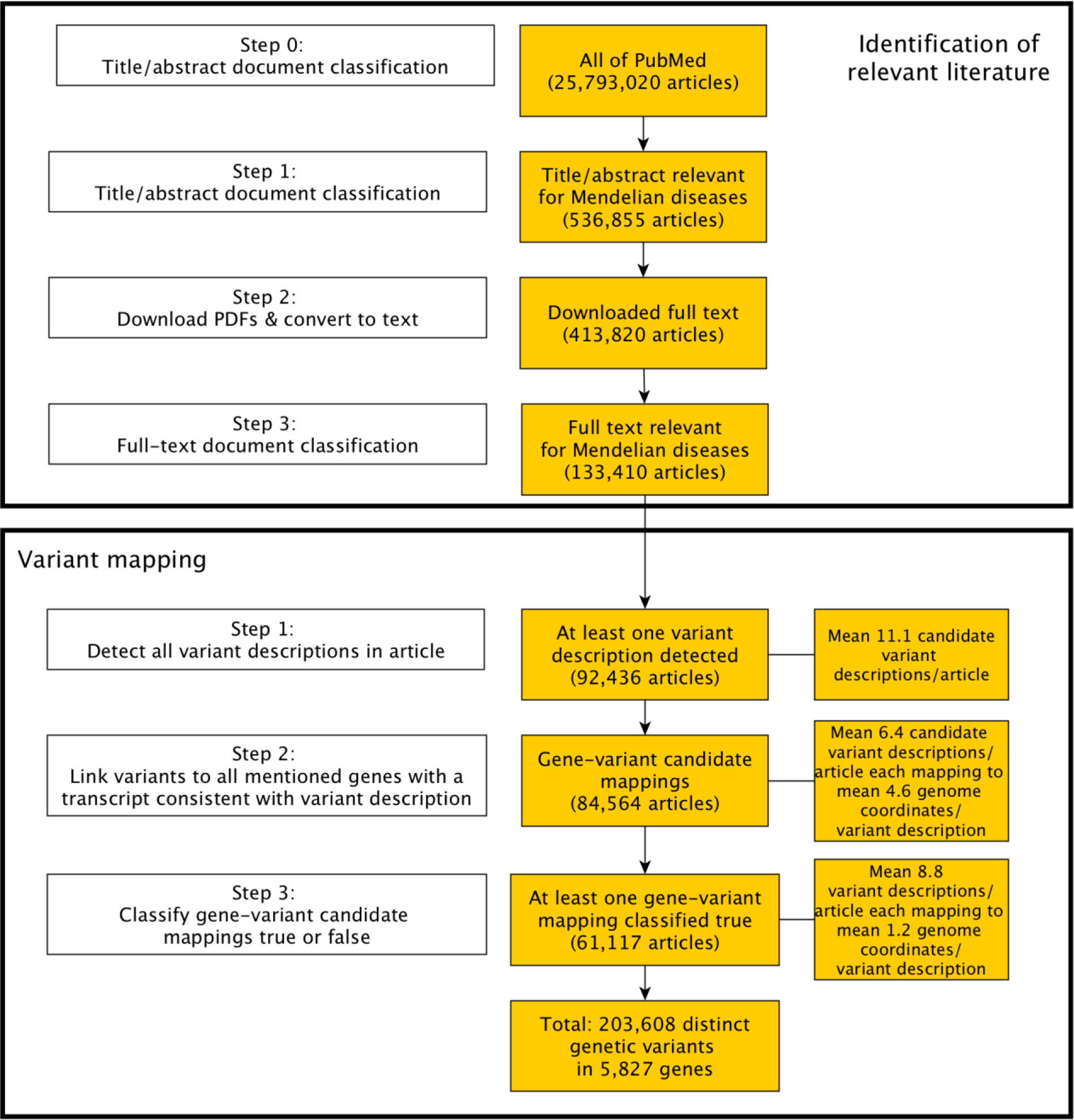
Construction of the automated variant database AVADA. Identification of relevant literature: Step 0: titles and abstracts of articles are downloaded from PubMed. Step 1: a suitable subset of relevant literature is identified by a document classifier that classifies titles and abstracts deposited in PubMed as possibly relevant or irrelevant to genetic disease. Step 2: full text PDFs of potentially relevant articles are downloaded wherever possible and converted to text. Step 3: the relevance of each paper to genetic disease gene knowledge extraction is reassessed using a full text document classifier. **Variant mapping**: Step 1: gene mentions are detected using a list of gene names and synonyms, and variant mentions are detected using 47 manually built regular expressions (Figure 2A). Step 2: a super-set of possible gene-variant candidate mappings is constructed out of all mentioned variants and genes in a paper where the variant appears to “fit” the gene: e.g., if a variant description is “c.123A>G”, the variant fits all genes mentioned in the paper that have at least one transcript with an “A” at coding position 123 (Figure 2B). Step 3: A machine learning classifier using a number of textual features (Figure 2C and Methods) describing the relationship between variant and gene mention in the article’s full text decides which of the previously constructed gene-variant candidate mappings are true, i.e., which variant actually refers to which gene (Figure 2D). AVADA extracts 203,608 distinct genetic variants in 5,827 genes from 61,117 articles.

### Variant and gene mention detection

In order to extract genetic variants from the full-text articles about human genetic disease and convert them to genomic coordinates, it is necessary to detect both mentions of genes and variant descriptions in articles about genetic disease. Extracting variant descriptions alone does not suffice, because variant descriptions in HGVS notation, such as “c.123A>G”, can only be converted to genomic coordinates if a transcript of the gene that the variant refers to is identified (Table 1).

**Table 1.** Examples of HGVS or common HGVS-like variant descriptions. Each row contains examples of a disease-causing variant description in HGVS or a common HGVS-like notation. Each of these variant descriptions describes a single genetic event causing a disease, usually by giving at least the position of the change in the gene’s transcript, an optional reference sequence and a novel alternative (mutated) sequence. All given variants can be described using multiple commonly used notations. Examples of alternatives to the notations are shown in the left hand column that denote the exact same genetic variants. Transcript identifiers for variant descriptions, which enable the mapping of variants to reference genome positions, are usually omitted by article authors, and must therefore be inferred by automated methods like AVADA. The right hand column lists the disease along with two articles using the variant descriptions given in the left hand column. The difficulty of parsing different variant notations that refer to the same genetic event warrants the development of automated approaches for variant curation from the literature.

| **HGVS(-like) variant descriptions (alternatives describing same genetic event)** | **Explanation of HGVS variant description** | **Disease caused by variant (cited literature uses all variant notations shown in left column)** |
| --- | --- | --- |
| NM_175073.2<br>593C>T<br>(NP_778243.1<br>p.A198V) | **DNA single nucleotide substitution**<br>reference C replaced by alternative T at position 593 in the transcript NM_175073.2 | Cerebellar ataxia with oculomotor apraxia type 1<sup>44,45</sup> |
| NM_006005.3<br>460+1G→A<br>(NM_006005.3<br>IVS4+1G>A) | **Splicing variant**<br>reference G replaced by alternative A at the genomic position 1 basepairs downstream of the 3' end of the exon of transcript NM_006005.3 that ends at position 460 | Wolfram syndrome<sup>46,47</sup> |
| NP_000518.1<br>p.Asp221Thrfs*44<br>(NM_000527.4<br>c.660delC;<br>NP_000518.1<br>p.Pro220Profsx45) | **Protein frameshift variant**<br>reference aspartic acid at residue number 221 in transcript NP_000518.1 impacted by an indel resulting in an alternative threonine, with the rest of the protein being frameshifted, introducing a stop codon 44 amino acid residues downstream of residue number 221 | Familial hypercholesterolaemia<sup>48,49</sup> |

AVADA extracts gene mentions from articles’ full text using a custom-built database of gene names containing gene name entries from the HUGO Gene Nomenclature Committee (HGNC) and UniProt databases. Gene and protein names from these were matched case-insensitive to word groups of length 1-8 in the document to identify gene mentions. To identify variant mentions, we manually developed a set of 47 regular expressions based on commonly observed HGVS-like variant notations in articles about human genetic disease (Supplementary Methods, Supplementary Table S1 and Figure 2A). At this step, we refer to every string that matches one of the 47 regular expressions as a “variant description”. In the AVADA full-text articles, variant descriptions in 92,436 articles were identified, with a mean of 11.1 variant descriptions per article (Figure 1).

**Figure 2.**
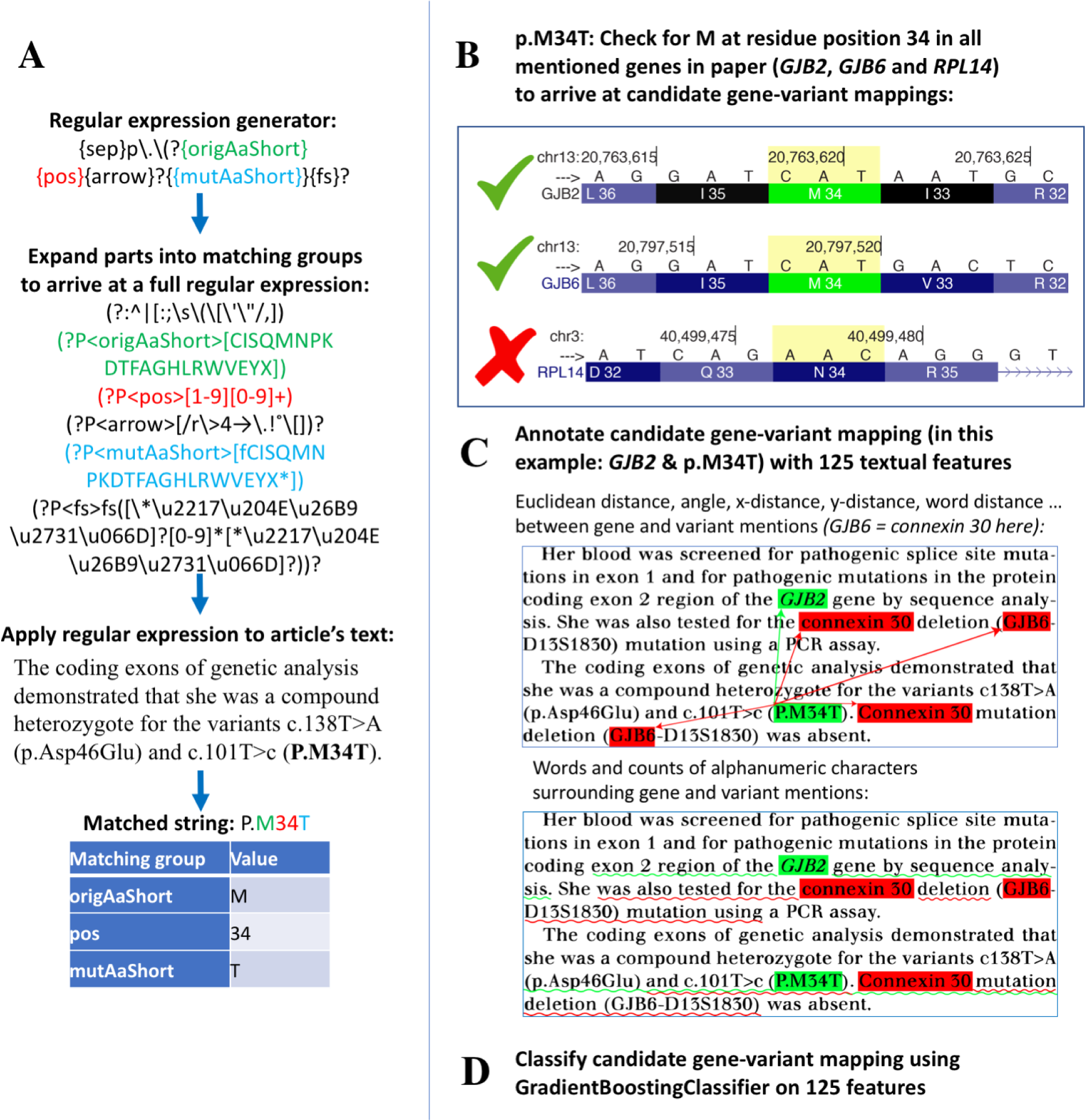
Automatic conversion of variant mentions to genomic coordinates from full-text literature. **(A)** AVADA uses 47 different regular expressions to detect variants in articles. Regular expressions are designed in forms of regular expression generators such as “{*sep}c\.{pos}{space}?{plusMinus}{space}?{offset}[,]*?{origDna}{space}?{arrow}{space}?{mutDna}*”. These regular expression generators contain named matching group generators, such as “*{origDna}*” (reference nucleotide, such as “A” or “T”) or “*{pos}*” (numeric position of the mutated nucleotide relative to the start of the transcript). Named matching group generators describe parts of the HGVS description that contain information about the variant. Regular expression generators are expanded into regular expressions by replacing the matching group generators, such as “*{pos}*”, into a named matching group, such as “*(?P<pos>[1-9][0-9]*)*”. Expanding all named matching group generators into named matching groups gives a full regular expression. If a full regular expression matches any string in a given article, the matched string is assumed to be a variant description. **(B)** Given a detected variant description and a set of genes detected in the text of an article, AVADA first checks if the variant matches any of the gene’s transcripts. In the current example, the variant p.M34T matches transcripts of the genes *GJB2* and *GJB6* because both have a methionine residue at position 34, but not the gene *RPL14* (with an asparagine at position 34). The variant p.M34T therefore forms gene-variant candidate mappings (p.M34T, *GJB2*) and (p.M34T, *GJB6*), which are filtered in the next step. **(C)** Given a gene-variant candidate mapping (variant=p.M34T and gene=*GJB2* in this example, highlighted in green), AVADA lets a Gradient Boosting classifier decide if the variant refers to the candidate gene using a set of 125 numerical features that contain information about the textual relationship between the variant mention and the textually closest mentions of the candidate gene (*GJB2*), as well as textually closest mentions of alternative nearby mentioned genes (connexin 30 (encoded by *GJB6*) in the example, in red). The 125 features are based on the relative positions of the closest candidate gene mentions to the variant mention, closest alternative gene mentions to the variant mention, information about the genes’ importance in the article, and words and characters surrounding the gene and variant mentions (see Methods). **(D)** The Gradient Boosting classifier takes these 125 features as input and returns a probability between 0 and 100% indicating the classifier’s assessment of whether the variant actually refers to the given candidate gene. If the classifier returns a likelihood greater than 90%, the gene-variant candidate mapping is transformed to Variant Call Format (chromosome, position, reference and alternative alleles) and entered into the AVADA database. In the present example, AVADA correctly decides that p.M34T only maps to *GJB2* and not connexin 30 (encoded by the gene *GJB6*). Example taken from PubMed ID 23808595.

### Mentioned genes form gene-variant candidate mappings with all mentioned variants that “fit” the gene

Having identified gene mentions and variant descriptions in text, it is now necessary to link variant descriptions with the genes that they refer to. Articles often mention variant descriptions without explicitly stating to which gene each variant description maps. The gene to which each variant description maps can be inferred by expert readers of the article. However, an automatic algorithm cannot easily infer to which gene a variant description maps, because gene mention and variant description do not necessarily occur in the same sentence or even the same paragraph or page.

To identify which variant description maps to which mentioned gene in the article, AVADA first forms so-called *gene-variant candidate mappings* between each variant description and each mentioned gene if the variant appears to “fit” at least one RefSeq^33^ transcript of the gene. Given an extracted variant description “c.123A>G”, the variant description forms gene-variant candidate mappings with all mentioned genes that have an “A” at coding position 123 of at least one transcript (Supplementary Methods and Figure 2B). A variant description can form gene-variant candidate mappings with multiple genes, which are filtered in the next step. Gene-variant candidate mappings are converted to genomic coordinates in the GRCh37/hg19 reference assembly. In the AVADA full-text articles, an extracted variant description initially mapped to a mean of 4.6 different genomic coordinates (Figure 1).

### Machine learning classifier selects the correct gene-variant mapping out of multiple gene-variant candidate mappings

AVADA uses a machine learning framework to decide which gene-variant candidate mappings are likely to be correct. The machine learning classifier is a scikit-learn GradientBoostingClassifier^34^. The training set for the classifier comprised positive gene-variant mappings curated from the literature in ClinVar, and a set of negative gene-variant mappings created by assigning variants from the positive training set to genes mentioned in the paper to which they did not map. Each gene-variant mapping was converted to a feature vector, based upon which the classifier decided if the gene-variant candidate mapping was true or false. The feature vector included the Euclidean distance between the 2D coordinates (consisting of page number, x and y coordinates of a mention) of the closest mentions of the variant and the gene in the PDF, the number of words between variant and gene mentions, the number of short “stopwords” (like “and”, “or”, “of”, …) around gene and variant mentions, and a number of other textual features containing information about the relationship between gene and variant mentions (Supplementary Methods and Figure 2C; performance analyzed below).

The classifier successfully reduced 4.6 candidate gene-variant mappings per variant description to a mean of 1.2 genomic positions in the AVADA full-text articles (Supplementary Methods and Figures 1, 2D).

## Results

### AVADA identified 203,608 variants in 5,827 genes from 61,117 articles

A total of 61,117 articles made it into the final AVADA database, with a mean of 8.8 identified variant descriptions per article. From these articles, 203,608 distinct genetic variants in 5,827 genes were automatically curated (Figure 1), comprising a variety of different variant types in a distribution strikingly similar to that of manually curated HGMD and ClinVar: for each of 6 categories of variant (stoploss, nonframeshift, splicing, stopgain, frameshift, missense), the fraction of variants AVADA extracted are between the fraction of the respective category in HGMD and ClinVar ±1% (Table 2). The articles used to construct AVADA are from a variety of journals, which are similar to the journals targeted by HGMD to curate its variants (9 out of the top 10 journals being the same between AVADA and HGMD; Figure 3A,B).

**Table 2.** Variant type percentages in AVADA, HGMD and ClinVar. Despite being based purely on automatic Natural Language Processing methods, AVADA variant type fractions are always within the range between manually curated HGMD and ClinVar ± 1%.

| **Variant type** | **AVADA** | **HGMD** | **ClinVar** |
| --- | --- | --- | --- |
| stoploss | 0.30% | 0.14% | 0.10% |
| nonframeshift | 2% | 3% | 3% |
| splicing | 8% | 7% | 4% |
| stopgain | 12% | 14% | 9% |
| frameshift | 14% | 22% | 11% |
| missense | 65% | 53% | 74% |

**Figure 3.**
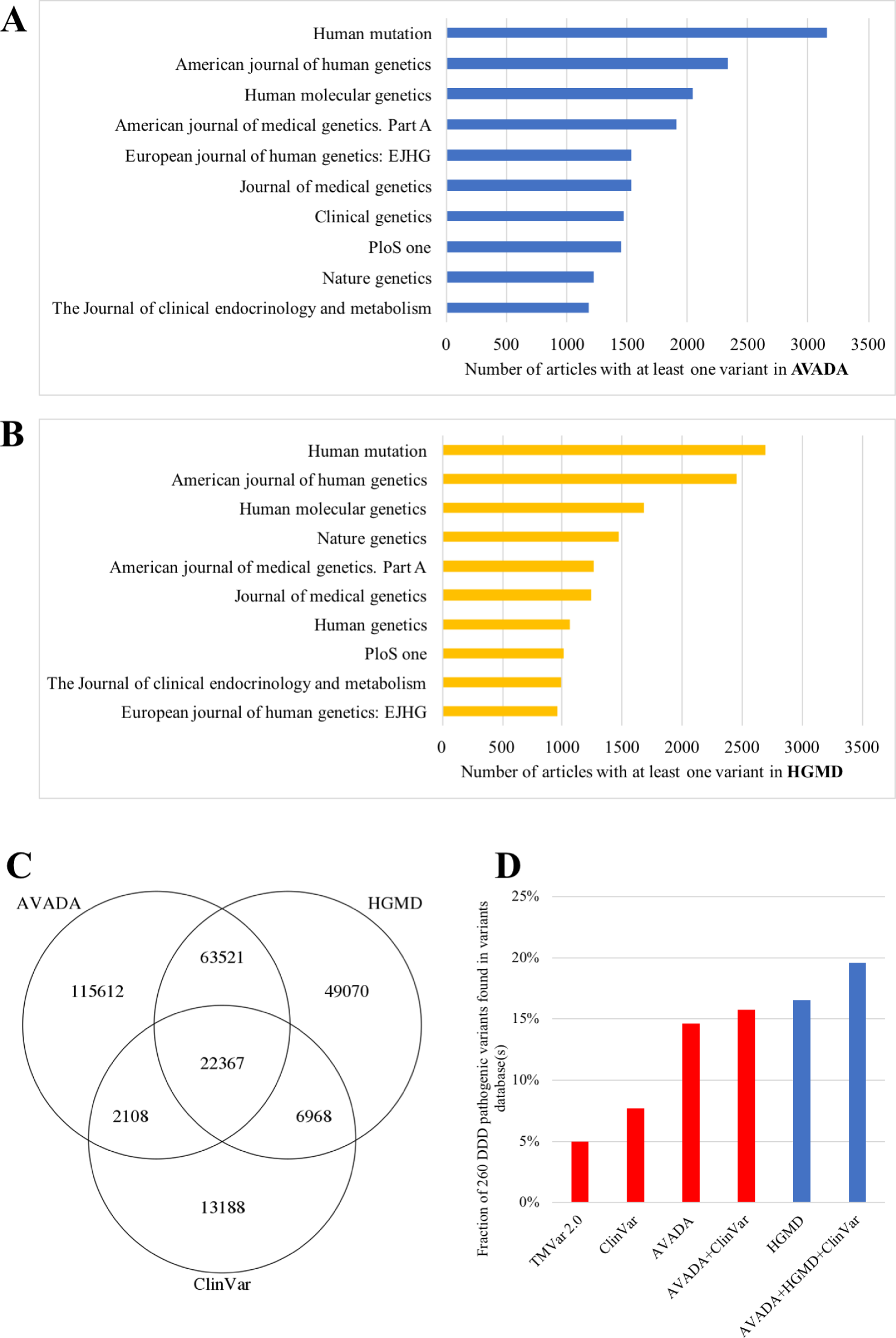
Automatic variant curation results. **(A)** Top ten journals in terms of number of articles curated in AVADA. AVADA extracted variants from 3,159 articles in “Human Mutation”, 2,330 articles in “American Journal of Human Genetics”, 2,042 articles in “Human Molecular Genetics” etc. **(B)** Top ten journals in terms of number of articles curated in HGMD. Similarly to AVADA, the top three journals are “Human Mutation”, the “American Journal of Human Genetics”, and “Human Molecular Genetics”. The two lists share 9 of the top 10 journals even though HGMD is manually curated whereas AVADA is entirely based on automated curation. **(C)** Extracted variants in AVADA intersected with all disease-causing variants in HGMD and ClinVar. AVADA extracts 85,888 variants in literature-based HGMD (subset to disease-causing variants) and 24,475 variants in submission-based ClinVar (subset to pathogenic and likely pathogenic variants). **(D)** Comparison of the fraction of Deciphering Developmental Disorders (DDD) causative variants found in various combinations of databases. 260 different variants were reported to be causative of 245 patients’ diseases in the DDD project, a large-scale diagnostic sequencing research project. We subset AVADA, HGMD, ClinVar and the automatically curated variant database tmVar 2.0 to sources pre-dating the publication of the DDD patient set. Of the causative variants, tmVar 2.0, which automatically parses on PubMed abstracts, contained 5%, ClinVar contained 8% reported as (likely) pathogenic, full text-based AVADA contained 15% and HGMD contained 17% reported as disease-causing. All tmVar 2.0 variants were either in AVADA or ClinVar. Combining the free (bars in red) AVADA and ClinVar databases recovers 16% of causative variants. Combining all databases facilitates rapid diagnosis for 20% of causative variants.

Each variant, defined by chromosome, position, reference and alternative allele, is annotated with: PubMed ID(s) of publications where this variant was extracted from; HUGO Gene Nomenclature Committee^35^ (HGNC) gene symbol, Ensembl ID^36^, and Entrez ID^37^ of the gene in which the variant is found, the inferred variant effect (e.g., “missense”), the RefSeq ID of the gene’s transcript to which the variant was mapped (e.g., NM_005101.3), and the exact variant description from the original article (e.g., “c.163C.T”). The latter allows clinicians to later rapidly locate mentions of this variant within the body of the article.

### AVADA is 72% precise

To estimate the precision (the fraction of extracted variants that are correctly extracted), 100 distinct random variants mapped to genomic coordinates by AVADA were manually examined. AVADA variants were manually counted as true extractions whenever the scientist reading the paper (using all lines of evidence in the paper such as Sanger sequencing reads, UCSC genome browser shots etc.) independently mapped the paper’s variant mention to the same genomic coordinates as AVADA. Of the 100 distinct random variants, 72% were extracted and mapped to the correct genomic position in GRCh37/hg19 coordinates without error by AVADA (Supplementary Table S2).

### AVADA recovers nearly 60% of disease-causing HGMD variants directly from the primary literature

We compared AVADA to HGMD and ClinVar versions with synchronized time stamps (Supplementary Methods). 85,888 AVADA variants coincided with variants marked as disease-causing (“DM”) in HGMD, corresponding to 61% of all disease-causing variants in HGMD. From this set of 85,888 AVADA variants, we selected 100 random variants and manually verified that the genomic coordinates (chromosome, position, reference and alternative alleles) were correctly extracted and the variant was reported as disease-causing in 97% of them (Supplementary Table S3). Thus, we infer that AVADA contains 59% of all disease-causing variants identified by HGMD.

We compared AVADA’s performance to the best previously published automatic variant curation tool, tmVar 2.0^27^, which attempts to map variant mentions in all PubMed abstracts to dbSNP identifiers (rsIDs). tmVar extracted only 19,424 disease-causing HGMD variants, or 14% of HGMD (Supplementary Figure 1 and Figure 3C). Considering only single nucleotide variants (SNVs), the largest class of known pathogenic variant, AVADA contains 70% of all DM SNVs in HGMD, of which an estimated 97% were extracted correctly. Similarly, AVADA contains 55% of all likely pathogenic or pathogenic variants in ClinVar (clinical significance level 4 or 5) and 62% of pathogenic or likely pathogenic SNVs in ClinVar. tmVar 2.0 extracted only 13,664, or 31%, of pathogenic or likely pathogenic variants in ClinVar.

Strikingly, AVADA contains 63,521 variants that are in HGMD (“DM” only) but not in ClinVar (clinical significance level 4 or 5). An analysis of a representative subset of 100 of the remaining 115,612 variants that were extracted by AVADA, but not reported as disease-causing in either HGMD or ClinVar, revealed them to be mostly benign or incorrectly extracted variants (Supplementary Table S4).

### Diagnosis of patients with Mendelian diseases using AVADA

We analyzed the accuracy of patient variant annotation with AVADA, tmVar, ClinVar and HGMD using a set of 245 patients from the Deciphering Developmental Disorders^38^ (DDD) study, harboring 260 causative variants reported by the original DDD study. De-identified DDD data were obtained from EGA^39^ study number EGAS00001000775 (Supplementary Methods). The DDD study is a large-scale sequencing study in which children affected with developmental disorders were sequenced in search of a molecular diagnosis. Disease-causing variants reported in DDD were obtained from Supplementary Table 4 in reference^38^.

### Sensitivity of variant annotation using AVADA, tmVar, HGMD and ClinVar

The more complete a variant database is, the higher its sensitivity when annotating patient genomes and the higher the likelihood of finding a causative variant in the patient’s genome. We determined how many of the 260 causative DDD variants were found in AVADA, tmVar, HGMD and ClinVar. The more causative variants are found in a database, the more rapidly some patients can be diagnosed. For the DDD patient variant annotation comparison, we subset AVADA and tmVar 2.0 to reference only articles until 2014 (before the publication of the DDD study), HGMD to use only variants added until 2014, and took the latest ClinVar version from 2014 (ClinVar version 20141202).

Of 260 different causative variants reported by the DDD study, a total of 45 variants were found by AVADA in the scientific literature. For each of these variants, all articles from which the variant was extracted were manually inspected. The variant was counted as correct if at least one article was found in which the variant’s genomic coordinates (chromosome, position, reference and alternative allele) were correctly extracted, the variant was reported as causative and the article did not cite the DDD study (pre-publication). 38 of the 45 variants found by AVADA fulfilled these criteria (Supplementary Table S5).

Only 20 variants reported to be causative by the DDD study were listed in ClinVar and ascribed a pathogenicity level of “pathogenic” or “likely pathogenic”. 43 variants were in HGMD, reported as “DM” (disease-causing). tmVar 2.0 contained 13 causative variants (Supplementary Table S6). AVADA and ClinVar together contained 41 causative variants. All of tmVar’s variants were either in AVADA or ClinVar. Thus, combining the free variant databases AVADA and ClinVar resulted in our annotating almost as many causative variants as are listed in HGMD. Combining all three databases yielded 51 variants (Figure 3D).

## Discussion

We present AVADA, an automated approach to constructing a highly penetrant variant database from full-text articles about human genetic diseases. AVADA automatically curated nearly a hundred thousand disease-causing variants from tens of thousands of downloaded and parsed full-text articles. All AVADA variants are stored in a Variant Call Format^40^ (VCF) file that includes the chromosome, position, reference and alternative alleles, variant strings as reported in the original article, and PubMed IDs of the original articles mentioning the variants. AVADA recovers nearly 60% of all disease-causing variants deposited in HGMD at a fraction of the cost of constructing a manually curated database^41^, over 4 times as many as the tmVar 2.0 database that relies on PubMed abstracts, and maps only to dbSNP rsIDs. From a cohort of 245 previously diagnosed patients from the Deciphering Developmental Disorders (DDD) project, AVADA pinpoints 38 DDD-reported disease-causing variants, fewer than HGMD (43) but almost twice as many as ClinVar (20) and almost three times as many as tmVar 2.0 (13), showing that this new resource will be useful in clinical practice. Combining the free variant databases AVADA and ClinVar recovers 41 diagnostic variants. This shows that AVADA is an important step into the direction of using machine learning approaches to improve the throughput of variant interpretation as proposed by ClinGen^26^.

Multiple lessons were learned from AVADA. First, curating variants from full text articles scattered between dozens of publishers’ web portals is worth the extra effort. However, while gene to variant linking is often relatively simple in the context of an abstract, this task is much more challenging in the context of sprawling full texts that may well discuss many additional genes beyond the causal few. A two-pronged approach is therefore necessary to further improve AVADA’s precision. First, our ability to link variants to the correct transcripts and genes can be improved. Second, non-pathogenic mentioned variants need to be better distinguished from pathogenic mentioned variants. Implementing patterns for more exotic variant notations and parsing supplements of articles would improve sensitivity, but also decrease precision.

AVADA curates variants without costly human input and can be re-run continually to discover newly reported variants without incurring significant additional cost. While the approach cannot currently replicate manual curation efforts, it is nevertheless well suited to support the work of manual curators in improving and extending existing variant databases. Blending the AVADA automatic variant curation approach with manual verification should facilitate rapid variant classification^42^ and the cost-effective annotation of patient variants.

Publishers can help further improve the automatic variant curation process by supplying database curation tools with simpler, stable programmatic access to full text and supplementary data of appropriate articles, a win-win step that would lead to both better variant databases, and increase the circulation of articles among their target audience. Requiring authors to abide by strict HGVS notation would also help. Moreover, the approach presented here can be extended to the automatic curation of genetic variants (in canonicalized representation) from other valuable modalities, such as somatic variants in cancer genes, animal models, cell lines, or non-model organisms with reference genomes and transcripts. The approach described could therefore support the rapid and cost-effective creation and upkeep of multiple different variant databases beyond human genetic diseases^43^ directly from the primary literature.

By comprehensively annotating each variant with information from the original articles (such as the originally reported variant string), AVADA enables rapid re-discovery and verification of a large fraction of reported variants in the scientific literature. Previously, manual curation efforts such as HGMD^24^ have demonstrated the power of systematic curation of pathogenic variants from the primary literature. AVADA shows that automatic variant curation from the full text literature is feasible and useful with regard to accelerating the creation of genetic variant databases. Combining automatic curation approaches like AVADA with manual curation will enable the creation and upkeep of cheaper, better, faster updating variant databases from the primary literature enabling both rapid diagnosis^42^ and reanalysis^12^.

## Acknowledgments

This work was funded in part by a Bio-X Stanford Interdisciplinary Graduate Fellowship to J.B.; by grants EMBO ALTF292-2011 and NIH/NHGRI 5U41HG002371-15 to M.H.; and by DARPA, the Stanford Pediatrics Department, a Packard Foundation Fellowship, a Microsoft Faculty Fellowship and the Stanford Data Science Initiative to G.B. We are obliged to thank the European Genome-Phenome Archive^39^ (EGA) and the Deciphering Developmental Diseases^38^ (DDD) project. The DDD study presents independent research commissioned by the Health Innovation Challenge Fund [grant number HICF-1009-003], a parallel funding partnership between the Wellcome Trust and the Department of Health, and the Wellcome Trust Sanger Institute [grant number WT098051]. The views expressed in this publication are those of the author(s) and not necessarily those of the Wellcome Trust or the Department of Health. The study has UK Research Ethics Committee approval (10/H0305/83, granted by the Cambridge South REC, and GEN/284/12 granted by the Republic of Ireland REC). Deidentified DDD data was obtained through EGA. The research team acknowledges the support of the National Institute for Health Research, through the Comprehensive Clinical Research Network.

## Author Contributions

J.B. and M.H. wrote software to map variants to the reference genome using a database of RefSeq transcripts. A.P.T. and J.B. verified AVADA-extracted variants. J.B. wrote the machine learning classifiers and performed performance evaluations. J.B., M.H., A.P.T., and G.B. wrote the manuscript. C.D. and K.A.J. downloaded and processed DDD data. P.D.S. and D.N.C. created HGMD and helped with manual variant inspection. J.A.B. provided guidance on clinical aspects of study design, testing set construction and interpretation of results. G.B. supervised the project. All authors read and commented on the manuscript.

## Web resources

All code for automatic variant curation with AVADA, as well as the automatically curated variants database presented here, will be available upon publication for non-commercial use at http://bejerano.stanford.edu/AVADA.

## Conflicts of interest

The authors declare no conflicts of interest.

## Supplementary Methods

### Variant Extraction Directly from Primary Literature

#### Download of literature

Articles were identified as potentially relevant based upon title and abstract in PubMed as previously described^9^. Briefly, all 25,793,020 available titles and abstracts from PubMed were downloaded. Subsequently, we trained a scikit-learn^28^ LogisticRegression^29^ classifier featurized by TF-IDF-transformed words (a common transformation of word frequencies into a feature vector). The training set for the title/abstract document classifier was based on 51,637 positive titles and abstracts cited in OMIM “Allelic Variants” sections or HGMD PRO version 2016.02, and 66,424 random negative titles and abstracts from PubMed. PDFs of articles were downloaded directly from publishers using PubMunch^50^.

#### Identification of relevant articles based on the full text of articles

We created a full-text classifier that assigns a score between 0 and 1 to each downloaded article, providing an estimate of the article’s likelihood of containing human pathogenic variant data. To create a TF-IDF feature vector, for use by a machine learning classifier, out of an article’s full text, each article was transformed by means of a scikit-learn CountVectorizer with parameters max_df=0.95 and min_df=100 followed by a TfidfTransformer with default parameters. The training set was based on 267,267 random articles in PubMed that were downloaded as a negative training set, and 46,291 full text articles cited in OMIM “Allelic Variants” sections or HGMD PRO version 2016.02. Based on this training set, a scikit-learn LogisticRegression^29^ classifier was trained.

#### Identifying candidate gene mentions in full text

Identification of candidate genes in full text was performed as previously described^9^. Briefly, a list of 188,975 gene and protein names was compiled from HGNC^35^ and UniProt^51^. Gene and protein names in this list were matched to word groups in the PDF text. Extractions were supplemented by PubTator^52^ gene extractions where available by matching gene names deposited in PubTator for a particular article to words occurring in that article.

#### Identifying candidate variant descriptions in full text

Candidate variant descriptions in Human Genome Variation Society (HGVS) or HGVS-like notation^53^ were identified using 47 regular expressions (Supplementary Table S1 and Supplementary Table S7). We partition mentioned variants into 3 broad categories: cDNA variants (“c.” variants, such as “c.123T>C”), protein variants (“p.” variants such as “p.T34Y”) and splicing variants (“c.” variants with a position and an offset, such as “c.123-2A>G” or “IVS” variants, such as “IVS4-2A>G”). Variant descriptions generally consist of a subset of the following components: variant type (cDNA, protein, splicing), position of the variant relative to the given transcript, reference nucleotide or amino acid, mutated nucleotide or amino acid, and type of genetic event (deletion, insertion, …). Using regular expression matching groups, information about all of these components is saved for each identified variant.

To create Figure 1, when counting the number of variant descriptions in articles, we removed all non-alphanumeric characters from variant descriptions because inconsistencies throughout the article with respect to spacing and parentheses used can otherwise lead to double-counting variant descriptions.

#### Mapping variants to candidate genes

A gene-variant candidate mapping of a variant onto a gene is a tuple *(g, v)* comprising a variant description *v* and a gene *g* such that there is at least one transcript *t* of *g* that has the variant’s given reference nucleotide/amino acid at the position given in the variant description *v.* If this is the case, the variant *v* is supported by the gene *g*, and *(g, v)* forms a candidate mapping.

To identify all gene-variant candidate mappings in an article with a set of mentioned variant descriptions *V* and a set of mentioned genes *G*, AVADA examines each pairwise combination *(g, v)* of a variant *v* in *V* and a gene *g* in *G* to determine if they form a candidate mapping. Each gene is represented by its set of transcripts deposited in the RefSeq^33^ database. All known RefSeq transcripts of *g* are successively examined to establish if *g* supports *v*. Most variants are written in a form that includes the position of the variant inside the gene’s transcript, the reference sequence, and the mutated sequence (e.g., “c.123A>G”: the position is “123”, the reference sequence is “A” and the mutated sequence is “G”). However, some variants only contain a position and a mutated sequence, not the original reference sequence (e.g., “c.153_154insGG”: the reference sequence is not included, just the novel insertion of “GG” between positions 153 and 154 inside the transcript). If the variant description *v* does not contain a reference sequence, all candidate genes form candidate gene-variant mappings with the variant. These gene-variant candidate mappings are further filtered using a machine learning classifier in the next section.

All gene-variant candidate mappings are converted to genomic coordinates (chromosome, position, reference allele and alternative allele). A conversion attempt is unsuccessful if the underlying nucleotide change cannot be identified given the variant description: e.g., this is the case for frameshift variants in “p.” notation such as “p.Val330fsX30”. Here, the precise underlying nucleotide change cannot be inferred from the variant description because the given frameshift may be caused by a very large number of possible nucleotide indel variants. In the case of a missense protein variant (e.g., NM_000025.2:p.Trp64Arg), the variant was translated to all possible single nucleotide variants that could cause such an amino acid change at the given position in the transcript. Since the Trp at position 64 in NM_000025.2 is encoded by the nucleotides TGG, both changing the T to a C (CGG) and the T to an A (AGG) result in an Arg codon. All further analysis was performed only on variants where conversion to genomic coordinates was successful.

#### Distinguishing true from false candidate gene-variant mappings

Given a set of candidate gene-variant mappings {(*g*_1_, *v*), (*g*_2_, *v*), (*g*_3_, *v*), (*g*_4_, *v*), …}, most of the genes *g*_i_ associated with *v* through a candidate mapping are false: the variant *v* does not map to gene *g*_i_. We constructed a machine learning classifier that distinguishes true gene-variant candidate mappings from false gene-variant candidate mappings. This classifier uses a vector of real numbers, called features, to determine if a gene-variant candidate mapping is true or false. In order to describe these features, some terminology must first be introduced:

- A “stopword” is a short word such as “by”, “of”, “there”, “if”, “or”, etc. The variant classifier uses a list of 122 stopwords (Supplementary Table S8).
- An alphanumeric character is a character in the ranges a-z, A-Z, and 0-9.
- A 2D position of a description in a PDF file consists of a page number and *x* and *y* coordinates of the mention on the page.
- A word position of a description in a PDF file consists of a single integer that gives the index of a word in the PDF document that contains the description.
- The Euclidean distance of two mentions associated with x and y coordinates (*x*_1_, *y*_1_) and (*x*_2_, *y*_2_) is defined as 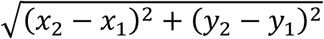
- The word distance between two mentions *m*_1_ and *m*_2_ of some genes or variants in an article *A* is defined as |*w*_2_-*w*_1_|.
- A mention *m*_1_ occurs “above” a mention *m*_2_ in the document if the page number of the 2D position of mention *m*_1_ is smaller than the page number of the 2D position of *m*_2_. If the page numbers of the two mentions are the same, *m*_1_ occurs before *m*_2_ if the y coordinate of *m*_1_ in the PDF is smaller than the y coordinate of *m*_2_ in the PDF.

Contextual information about a gene or variant mention in a PDF file is defined to consist of the following:

- the number of stopwords among the 20 words preceding the mention in the article’s text
- the number of stopwords among the 20 words following the mention in the article’s text
- the number of alphanumeric characters among the 20 characters preceding the mention in the article’s text
- the number of alphanumeric characters among the 20 characters following the mention in the article’s text.

Each gene *g* is mentioned 1 to *n* times in an article. Let *mention(g)*_1_ … *mention(g)*_n_ be the mentions of the gene *g* in the article. Similarly, each variant *v* is mentioned 1 to *m* times in an article. Let *mention(v)*_1_ … *mention(v)*_m_ be the mentions of the variant *v* in the article.

The machine learning classifier used by AVADA to distinguish true from false gene-variant candidate mappings is a scikit-learn GradientBoostingClassifier^34^. To decide whether a given gene-variant candidate mapping is true or false, the GradientBoostingClassifier takes a list of 125 numerical features containing information about the relationship between mentions of the gene and mentions of the variant in the original article. Based on these features, the classifier returns a number between 0 and 1 that gives the likelihood of the gene-variant mapping being true or not. The 125 features are constructed in 8 different feature groups describing the textual and geometric relationship between the candidate gene and candidate variant mention, and other genes mentioned close to the candidate variant mention. Further information is available in the accompanying code (see “variant_classifier_features.py”, functions “relationship_2d” and “relationship_wordspace”).

The variant classifier decides if *mention(v)*_j_ maps to gene *g* for 1 ≤ *j* ≤ *m* based on these 125 features. The value of these features is determined separately for each variant mention *mention(v)*_j_. If the classifier decides that any variant mention in *mention(v)*_1_ … *mention(v)*_m_ maps to *g* with classifier score greater or equal to 0.9, the variant *v* is considered to map to the gene *g*. To train the classifier, it was presented with a large number of annotated true and false gene-variant candidate mappings, called a training set. The training set for the classifier was created as follows: gene-variant candidate mappings *(g, v)* discovered by AVADA in a given article *A* were converted to genomic coordinates in form of chromosome, position, reference and alternative allele. If the genomic coordinates of a gene-variant candidate mapping extracted from *A* were deposited in ClinVar version 20170228 and annotated as curated from *A*, the mapping *(g, v)* was supervised true and all mappings of other genes to the same variant *v* in the article were supervised false. Otherwise, the variant was discarded. Synonymous variants (e.g., “p.Trp88Trp”) were also discarded due to the fact that they were largely not disease-causing, or were false extractions. This strategy yielded a training set comprising 25,218 positive training examples and 91,742 negative training examples from 7,823 articles. The importance assigned to each of the 125 features by the GradientBoostingClassifier is listed in Supplementary Table S9.

All extracted variants in AVADA were pre-processed using bcftools^54^ to normalize all variants (left-align indels and exclude variants where the RefSeq reference nucleotide did not match the GRCh37/hg19 nucleotide):

~~~
bcftools norm ––check-ref x-f human_g1k_v37.fasta-o avada.vcfavada_non_normalized.vcf
~~~

#### Comparison of AVADA to HGMD, ClinVar, and tmVar 2.0

The first version of AVADA was created on articles downloaded until June 2016. To ensure a fair comparison, we compare AVADA with HGMD PRO version 2016.02 and ClinVar version 20160705. These were obtained from www.hgmd.cf.ac.uk/ac/index.php and ftp://ftp.ncbi.nlm.nih.gov/pub/clinvar/vcf_GRCh37/, respectively. tmVar 2.0 variants were obtained from ftp://ftp.ncbi.nlm.nih.gov/pub/lu/PubTator/mutation2pubtator.gz. The tmVar file was subset to contain only tmVar-extracted variants in articles from 2016 and before (same set of articles used as input to AVADA). tmVar-extracted rsIDs were converted to genome coordinates by joining with the official dbSNP database mapping rsIDs to genome coordinates at ftp://ftp.ncbi.nlm.nih.gov/snp/organisms/human_9606_b151_GRCh37p13/VCF/All_20180423.vcf.gz.

Variants reported in AVADA, HGMD, ClinVar, and tmVar 2.0 were normalized (as above) using bcftools:

~~~
bcftools norm ––check-ref x-f human_g1k_v37.fasta-o<database_normalized>.vcf <database>.vcf
~~~

Variants were counted to be in two variant databases if the full variant description (chromosome, position, reference and alternative alleles) in both databases matched exactly. HGMD contained 165,051 distinct variants, of which 141,926 were marked as disease-causing (“DM”). ClinVar contained 142,396 distinct variants, of which 44,631 were marked as “pathogenic” or “likely pathogenic”. tmVar 2.0 contained 80,159 distinct variants.

#### Variant types contained in AVADA

To count the fractions of variant types contained in AVADA, each variant was assigned one of the types “missense” (single nucleotide variants changing an amino acid in the mapped gene), “nonframeshift” (insertion, deletion and indel variants adding a multiple of 3 nucleotides to a coding exon), “frameshift” (all other insertion, deletion and indel variants in coding exons), “splicing” (splice-site variants), “stopgain” (single nucleotide variants changing an amino acid codon in a coding exon to a stop codon) and “stoploss” (single nucleotide variants changing a stop codon to an amino acid codon) by automatically analyzing the effect of the variant on the mapped transcript. Variants of all types were summed, and fractions of variant types were calculated as the number of variants of a particular type over the total number of variants of all types in AVADA.

#### Variant types contained in ClinVar and HGMD

To generate fractions of variant types in HGMD and ClinVar, variants in these databases were annotated with semantic effect using ANNOVAR^55^. All HGMD or ClinVar variants that had a missense, stoploss, stopgain, splice-site, frameshift or nonframeshift effect in ENSEMBL^36^ and RefSeq^33^ coding exons, and had a variant frequency of less than 3% in ExAC^56^ v0.3 and the 1000 Genomes Project^57^ phase 3 were counted, and percentages of each variant type were calculated as the number of variants of a particular type over the total number of missense, stoploss, stopgain, splice-site, frameshift and nonframeshift variants in HGMD and ClinVar, respectively.

#### Diagnosis of patients with Mendelian diseases using AVADA

DDD patient Variant Call Format (VCF) files were obtained from the European Genome-Phenome Archive^39^ (EGA) study number EGAS00001000775. We identified VCF files for affected patients by matching the phenotypes that each VCF file was annotated with the phenotypes that each patient identifier and causative variant were annotated with, and verifying that the causative variant was contained in the patient’s associated VCF file. If unique identification of a patient’s VCF file was not possible, we omitted the patient. Reported disease-causing variants that were not found in a VCF file were omitted. Bcftools were used to normalize all variants in DDD VCF files using the following command:

~~~
bcftools norm-f human_g1k_v37.fasta-o <normed DDD VCF file><original DDD VCF file>
~~~

#### Sensitivity of variant annotation using AVADA, tmVar, HGMD, and ClinVar

ANNOVAR^55^ was used to annotate variants with a predicted effect on protein-coding genes from ENSEMBL^36^ and RefSeq^33^, and allele frequencies from the ExAC^56^ v0.3, the 1000 Genomes Project^57^ phase 3 and the UK10K^58^ ALSPAC and TWINS sub-cohorts. All variants with a frequency of at most 0.5% in all sub-populations of ExAC v0.3, 1000 Genomes Project and the UK10K ALSPAC and TWINS sub-cohorts, that affected a protein-coding gene and were missense, stopgain, stoploss, frameshift indel, nonframeshift indel or splice-site disrupting were retained.

AVADA and tmVar 2.0 were subset to variants from articles until 2014 by associating each article with the publication date stored in PubMed and subsetting to articles until 2014. HGMD variants were subset to 2014 by removing all variants with a “new_date” greater than 2014. ClinVar version 20141202 was obtained from ftp://ftp.ncbi.nlm.nih.gov/pub/clinvar/vcf_GRCh37/archive_1.0/2014/.

**Supplementary Figure 1.**
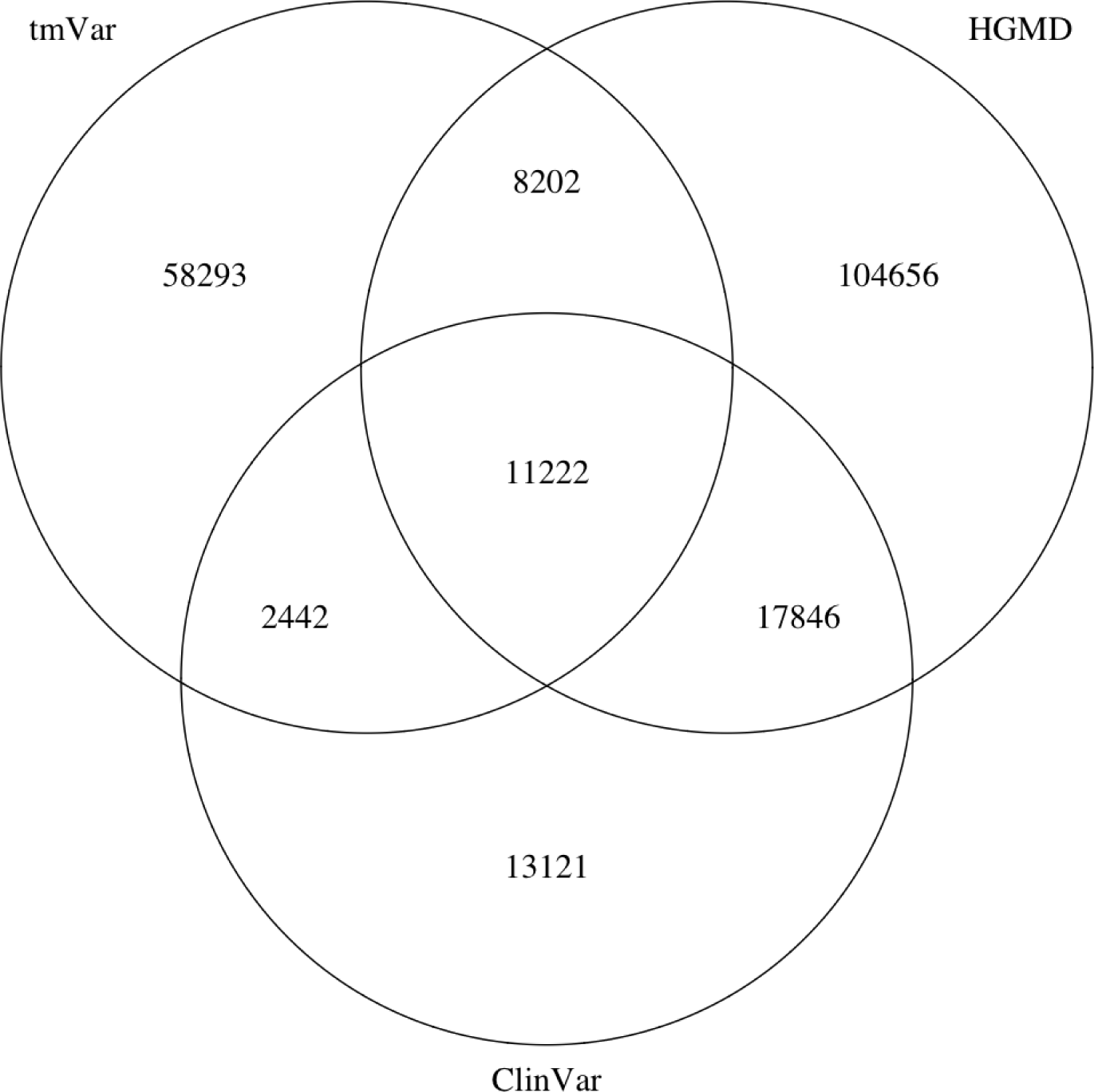
Extracted variants in tmVar intersected with all disease-causing variants in HGMD and ClinVar. tmVar extracts 19,424 variants in HGMD (subset to disease-causing variants), as compared to 85,888 variants for AVADA and 13,664 variants in ClinVar (subset to pathogenic and likely pathogenic variants), as compared to 24,475 for AVADA.

